# Timing appearance and integration of actin-organizing palladin protein in dynamic myofibril assembly

**DOI:** 10.1101/047183

**Authors:** Ngoc-Uyen-Nhi Nguyen, Tz-Yu Liu, Hao-Ven Wang

## Abstract

The involvement of actin-associated protein palladin in myogenesis has been elucidated, however, palladin distribution in a functional myotube remains to be identified. Since actin is required for myofibrillogenesis, it is of great interest to enhance our understanding of the spatial arrangements of palladin during sarcomeric assembly. Surprisingly, palladin was found to be discretely organized in different stages of myofibrillogenesis. Palladin revealed stress-fiber-like structures at undifferentiated stages, subsequently displayed chaotic expression and strongly co-distributed with actin, α-actinin, and myosin heavy chain of premyofibrils. At late stages, aggregates of palladin were spaced in a regular dot-like structure. On the other hand, palladin presents at I-Z-I bands of adult muscle. These observations suggest that palladin engages with sarcomeric proteins during the process of myoblast differentiation and that these interactions might occur in a temporally regulated fashion. In addition, transient overexpression of 140-kDa-palladin resulted in nonfilamentous actin arresting mature myotube formation. 200-kDa-palladin overexpression led to the early formation of Z-lines. Collectively, these findings suggest that palladin might serve a role in myofibrillogenesis by guiding and positioning sarcomeric proteins at the appropriate time and place. Our results highlight the involvement of palladin protein and the discrete functions of palladin isoforms in sarcomeric development in vitro.

## Introduction

Skeletal muscle differentiation is a multistep progress, in which small mononucleated myoblasts differentiate and progressively fuse to form larger multinucleated syncytia, called myotubes (Bentzinger et al., 2012; Dias et al., 1994; Sabourin and Rudnicki, 2000). Myotubes consist of muscle-specific proteins, assembling functional contractile units named sarcomeres (Charge and Rudnicki, 2004; Myhre and Pilgrim, 2012; Sanger et al., 2009). Sarcomeres are composed of actin (thin) and myosin (thick) filaments, and their associated proteins, making up the I-bands and A-bands, respectively. The tandem arrays of sarcomeres organize cylindrical organelles called myofibrils, which are responsible for precise muscle contraction.

The assembly of myofibrils has been proposed to develop from premyofibrils to nascent and subsequent mature myofibrils (Sanger et al., 2010; Crawford and Horowits, 2011). At the beginning of myofibrillogenesis, the primary component of Z-lines, α-actinin, appears as Z-bodies along stress-fiber-like structures (Rhee et al., 1994). The mini A-band non-muscle myosin II interdigitates the overlapping actin filaments and presents in between the Z-bodies, forming minisarcomeres of premyofibrils (Sanger et al., 2005). Subsequently, titin links up Z-bodies and myomesin, building a scaffold for thick filament integration into the sarcomeres. Thin filament proteins, such as nebulin, tropomodulin, and tropomyosin, are recruited to the actin filaments, and the premyofibrils turn into nascent myofibrils. The muscle myosin represented by four myosin light chains (MLCs) and two myosin heavy chains (MyHCs) replaces non-muscle myosin, attaching to the M-lines localized in the center of the A-bands. The sarcomeres of nascent myofibrils then grow in length from small Z-bodies to larger Z-lines, forming the center of the I-bands of mature myofibrils (Sanger et al., 2002; Nwe et al., 1999; Rui et al., 2010).

During differentiation, fused myocytes rearrange a non-muscle-specific cytoskeleton into a muscle-specific, contractive sarcomeric cytoskeleton, reflecting the vast structural reorganization of the actin cytoskeleton. Since contraction is actin-based, actin filaments and a subset of specialized cytoskeletal proteins are arguably the major functional cytoskeletal components in muscle. Palladin is an actin-remodelling protein known to co-distribute with numerous sarcomeric proteins such as actin, α-actinin, nebulin, and vinculin, suggesting the presence of palladin in myofibrillogenesis (Parast and Otey, 2000; Mykkanen et al., 2001). However, the precise distribution of palladin during sarcomere formation is not well known.

A single palladin gene yields various isoforms as a result of alternative RNA splicing and the usage of different promoters. Three canonical isoforms of palladin, with molecular weights of 200, 140, and 90-kDa, respectively, have been characterized in numerous studies. Recently, we have examined the involvement of palladin during skeletal muscle differentiation (Nguyen et al., 2014; Nguyen and Wang, 2015). Interestingly, Wang and Moser observed the localization of the largest isoform of palladin to the Z-discs of sarcomeres in overexpression experiments, both in primary cardiomyocytes and myotubes (Wang and Moser, 2008). Given that a sarcomere is the basic contractile unit capable of differentiation, the present study aims to clarify further the subcellular localization of palladin in sarcomeric formation in vitro by using the murine C2C12 skeletal muscle cell line.

## Materials and methods

### Antibodies and reagents

Dulbecco’s modified Eagle’s medium (DMEM) was obtained from Hyclone, USA. Penicillin-streptomycin was obtained from Nutricell, USA. Fetal bovine and horse sera were obtained from Gibco, USA.

The following reagents were used: G418 (Sigma-Aldrich, USA), Lipofectamine 3000 (Invitrogen, USA), Bio-Rad Protein Assay Dye Reagent (Bio-Rad Laboratories, USA), a protease and phosphatase inhibitor cocktail (Sigma-Aldrich), RIPA buffer (Cell Signaling Technology, USA), ECL immunoblotting kit (GE Healthcare, USA), and ProLong Gold Antifade Mountant with DAPI (Invitrogen). Horseradish peroxidase-conjugated secondary antibodies were purchased from Jackson ImmunoResearch, USA and used at a ratio of 1:5000.

The following primary antibodies were used: rabbit anti-palladin (1:400, ProteinTech, USA); mouse anti-MyHC (clone MY-32) (1:800, Genetex, USA); mouse anti-MLC (1:400, Genetex); mouse anti-α-actinin (1:500, Genetex). The following immunofluorescence secondary antibodies were obtained from Invitrogen and used at 1:400: Alexa Fluor-488 goat anti-rabbit IgG and Alexa Fluor-568 goat anti-mouse IgG. TRITC-phalloidin was purchased from Life Technologies and used at 1:400.

### Murine skeletal muscle cell line C2C12 culture

C2C12 cells were obtained from the Bioresource Collection and Research Center, Taiwan. Myoblasts were propagated in growth medium (GM) containing DMEM supplemented with heat-inactivated 10% fetal bovine serum, and 1% penicillin-streptomycin in a 5% CO_2_ atmosphere at 37C. For differentiation into myotubes, confluent C2C12 myoblasts were switched to a differentiation medium (DM) containing DMEM with 2% horse serum. The myotubes began to form 2-4 days post-differentiation. Every two days, the myotubes were fed with fresh DM. The cells were observed at two time points: at undifferentiated (myoblasts) and differentiated stages (myotubes).

### Western blotting

C2C12 cell cultures were collected at indicated time points, and protein was extracted using RIPA buffer with a protease-phosphatase inhibitor cocktail. The protein was quantified using the Bradford method. Equal amounts (50 μg) of protein extracts were resolved in SDS-PAGE, eletrophoretically transferred to polyvinylidene difluoride (PVDF) membranes, and subjected to immunoblotting. Membranes were blocked with 5% skim milk for 1 h at room temperature. Membranes were then incubated with primary antibodies overnight at 4C and with secondary antibodies for 1 h at room temperature.

### Transient transfection

Full length murine palladin isoforms were cloned into plasmid pEGFP-N1 (Clontech, Japan), yielding EGFP-90-kDa palladin, EGFP-140-kDa palladin, and EGFP-200-kDa palladin. C2C12 myoblasts were transfected with 5 μg of the linear plasmid and Lipofectamine 3000 reagent according to the manufacturer’s instructions. At 48 h post-transfection, the transfected-myoblasts were selected under 2 mg/mL of geneticin (G418) for an additional five days. Selected cells were seeded in equal numbers for one day in GM, and incubated for an additional 4 days in DM.

### Immunofluorescence

For mouse skeletal muscle sections, mice were sacrificed following the guidelines of the Affidavit of Approval of Animal Use Protocol of National Cheng Kung University (animal ethics number 103149). The hind limb muscle was dissected, placed in OCT blocks, and cryosectioned at −20C in 10 μm longitudinal sections. The sections were placed onto Superfrost Plus glass slides. For cell line sections, C2C12 cell cultures were grown on 22 mm × 22 mm glass cover slips in GM to 80% confluence and then switched to DM and harvested at indicated time points. Sections were fixed with 4% paraformaldehyde for 15 min, and permeabilized with 0.2% (v/v) Triton X-100 for 15 min. Non-specific binding of secondary antibodies was blocked by 1 h of incubation in 5% (w/v) bovine serum albumin in phosphate-buffered saline (PBS). Subsequently, sections were incubated at 4C overnight with primary antibodies diluted in a mixture containing 1% BSA, 0.1 % gelatin, 0.05% Tween20, and 0.01 M PBS. The following day, sections were washed with PBS, and then incubated with fluorochrome-conjugated secondary antibodies at room temperature for 1 h. Actin was stained with TRITC-phalloidin for 30 min. Immunofluorescence sections were mounted on slides with ProLong Gold mounting solution and visualized using a Nikon TE2000 Eclipse C1si confocal microscope and EclipseTi epifluorescence microscope (Nikon Inc., Japan). Images were processed with NIS-element software.

## Results

The association of palladin protein during myofibril assembly in myotubes differentiated from C2C12 myoblasts was examined. Cells were maintained in the proliferation state or induced to differentiate for 2 to 4 days to form myotubes. Samples were then processed for immunolabelling with polyclonal antibodies against the C-terminus of palladin and major myofibril proteins, including actin, α-actinin, and MyHC.

### Palladin expression pattern in cultured myotube formation

The palladin expression pattern in the differentiation of C2C12 cells was initially confirmed using Western blotting (Fig. 1A). In response to differentiation induction, the expression of muscle-specific factors, including myogenin, MyHC, and MLC, increased. The 200-kDa palladin protein level was gradually upregulated compared to that of day 0 cultures while 140-kDa palladin expression slightly decreased following differentiation. 90-kDa palladin protein was found in myoblasts, and increased during the differentiation process. By day 3 of differentiation, most myoblasts (Fig. 1B) became elongated and spindle-shaped (Fig. 1B’, arrows). The developing myotubes appeared strongly actin-(Fig. 1D), α-actinin-(Fig. 1E), and MyHC-positive (Fig. 1F), and thus could be promptly distinguished from the surrounding mononucleated cells. Similar to these sarcomeric proteins, palladin in myotubes was labelled with stronger signal than palladin in undifferentiated cells (Fig. 1C). In addition, double-immunofluorescence staining of C2C12 myotubes with MLC and α-actinin antibodies displayed a striation pattern of the sarcomeric cytoarchitecture (Fig. S1, see inset in C). Taken together, the C2C12 cell line matured sufficiently to form sarcomeric structures, which can serve as a simplified system for studying the temporal-spatial distribution of palladin during sarcomere assembly.

**Figure 1:**
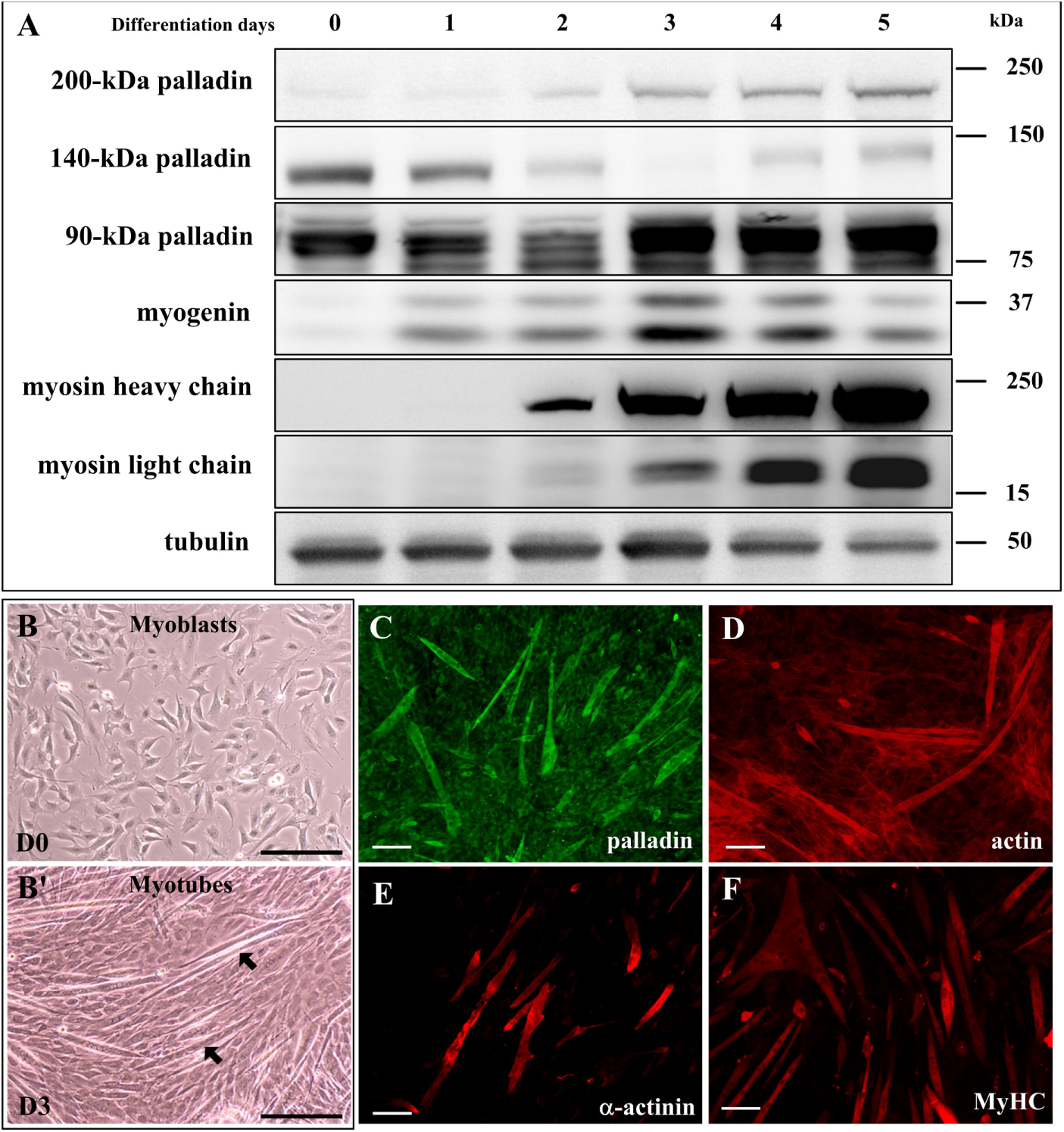
Palladin expression pattern during differentiation of C2C12 cells. (A) Western blot analyses were performed to determine endogenous palladin protein levels in C2C12 differentiated for 0-5 days. Myogenin and muscle myosins served as differentiation markers. Tubulin was used as a loading control. (B) Bright-field images of C2C12 myoblasts and (B’) myotubes were taken at 0 and 3 days of differentiation. Scale bar: 50 μm. (C-F) Immunofluorescence detection of (C) palladin, (D) F-actin, (E) sarcomeric α-actinin, and (F) sarcomeric MyHC in C2C12 cell cultures at day 3 of differentiation. Scale bar: 100 μm.

### Palladin co-localizes to actin, α-actinin and MyHC during nascent myofibrillogenesis in vitro

In the undifferentiated condition, the C2C12 myoblasts appeared mononucleated, flat, and fusiform. These myoblasts express continuous and defined filamentous actin that formed a randomly oriented network of filaments (Figs. 2A and S2A). Some myoblasts were lightly stained with α-actinin but were mainly concentrated in the perinuclear area (Figs. 2D and S3A, arrow). At this stage, myoblasts did not express relevant amounts of myogenic proteins. The expression of MyHC, which has been linked to advanced myogenic differentiation, could not be detected (Figs. 2G and S4A). Palladin proteins were widely identified and distributed diffusely in the myoplasm, where they co-localized as regularly-spaced dots along cytoplasmic actin stress fibers (Figs. 2A’ and S2C, white arrows), and also presented at the ends of filaments that surrounded focal adhesion sites (Fig. S2C, red arrows). Interestingly, palladin also accumulated around the nucleus (Figs. 2A’, 2D’, S2B-S4B), where it co-localized with α-actinin (Figs. 2D” and S3C).

**Figure 2:**
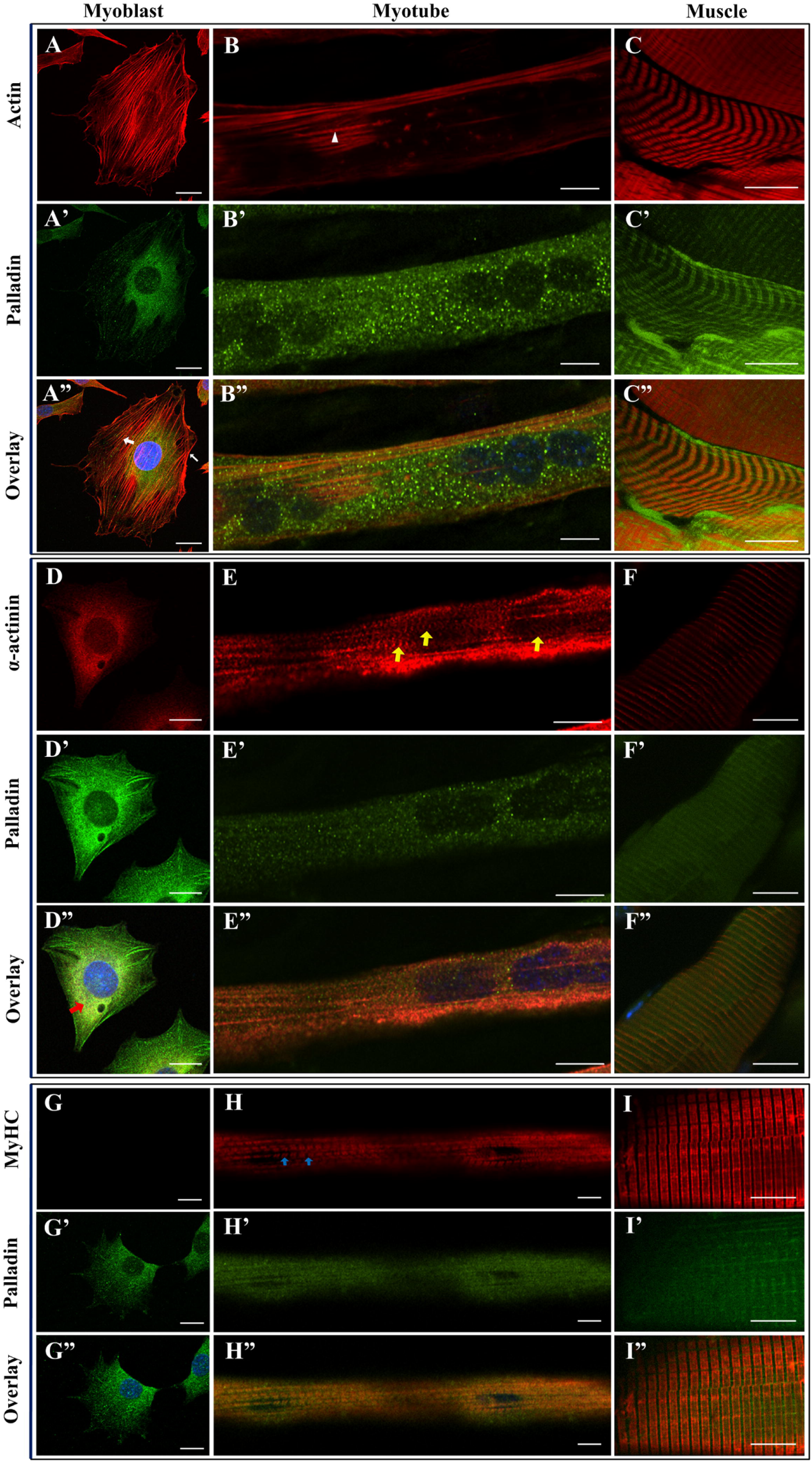
Localization of endogenous palladin in cultured myofibril assembly and adult skeletal muscle fibers. Panels depict C2C12 myoblasts (A, D, and G), myotubes (B, E, and H), and longitudinal cryosections of mouse hindlimb muscle immunostained for phalloidin-stained F-actin (A-C, red), actinin (D-F, red), MyHC (G-I, red), and endogenous palladin (green), and imaged by confocal microscopy. DAPI-stained nucleus is shown in blue. Left panels: in myoblasts, palladin colocalizes with actin at stress-fibers-like filaments (A”, white arrows), and sarcomeric α-actinin in the perinuclear area (D”, red arrow). Middle panels: during myofibril assembly, actin is reorganized into continuous, striated actin filaments (B, white arrowhead). Developing myofibrils also present a striated α-actinin distribution (E, yellow arrows) and a sarcomere-like periodic pattern of MyHC (H, blue arrows) after 4 days of differentiation. Palladin associates with acin, α-actinin, and MyHC during in vitro myofibril formation. Palladin is distributed as dot-like structures between myofibril filaments. Right panels: confocal sections of mouse hindlimb skeletal muscle show the concentration of palladin in the I-Z-I bands. Note that palladin is enriched at the Z-line.

Two days after differentiation, some nascent myotubes that resulted from the fusion of myoblasts were detected. They mainly existed in an elongated shape with a spatial organization of premyofibrils (Figs. S2F-4F). Their actin filaments were remodeled into long and continuous filaments than expanded along the longitudinal axis of the myotubes (Fig. S2D). Sarcomeric α-actinin aggregated into a punctate pattern along stress-fiber-like structures that resembled Z-bodies, preferentially presented at the myotube periphery (Fig. S3D, arrowhead). In addition, MyHC began to upregulate during this stage (Fig. S4D). In the nascent myotubes, palladin was extensively distributed throughout the cytoplasm (Figs. S2E-S4E). It is noteworthy that palladin co-localized with not only actin (Fig. S2F) and α-actinin (Fig. S3F), but also MyHC (Fig. S4F), in the premyofibril stages.

### Palladin mainly contributes to I-Z-I bands in mature myotubes

At day 4 of differentiation, multinucleated myotubes had numerous myonuclei that were centrally localized, clumped, or distributed all along the longitudinal direction of myotubes (Figs. 2B, E, H) and displayed the striated banding of myofibrils. Longitudinal actin filaments presented along the myotubes to form thin filaments with striated I-bands (Fig. 2B, white arrowhead). Sarcomeric α-actinin on the Z-bodies laterally coalesced to form the sarcomeric Z-lines (Fig. 2E, yellow arrows). MyHC was abundantly expressed and concentrated in the cytoplasm with the typical striated pattern of the A-bands (Figs. 2H and S4G, blue arrows). Palladin was also amply expressed in multinucleated myotubes, incorporated into myofibrils, and either continuously or occasionally showed primitive striations (Fig. S3H, yellow arrowheads). Merging images still revealed the co-localization of palladin with actin (Figs. 2B”, S2I) and sarcomeric α-actinin (Figs. 2E”, S3I, see inset). Since MyHC formed the striated structure (Fig. 2H, blue arrows) in this stage, palladin diffusely appeared in the cytoplasm arround MyHC bands (Fig. 2H”). Thus, palladin may associate with materials present between striated bands.

In adult hind limb skeletal muscle, palladin fluorescence signals were observed throughout the I-band region of muscle sarcomeres (Fig. 2C”), significantly enriched with muscle-specific α-actinin staining in the Z-line location (Fig. 2F”). In addition, palladin was found to be concentrated in the gap between MyHC bands (Figs. 2I” and S5). Thus, although palladin and MyHC formed preassembled complexes in the premyofibril, they seem to not co-localize in fully differentiated adult skeletal muscle fibers.

### Overexpressed palladin isoforms and Z-line formation

We have demonstrated that palladin co-localizes in the I-Z-I band with sarcomeric α-actinin and actin filaments. It is broadly accepted that palladin protein is composed of many isoforms. To further investigate the distribution of each isoform in myofibril assembly, C2C12 cells transfected with constructs encoding EGFP-tagged 90- (Figs. 3A and 3D), 140-(Figs 3B and 3G), and 200-kDa palladin (Figs 3C and 3J) were fixed and stained with sarcomeric α-actinin (Fig. 3) or actin (Fig. S6). It is noteworthy that the expression of recombinant palladin isoforms in C2C12 cells resulted in strikingly different patterns. EGFP-90-kDa palladin was expressed in a striated, closely spaced spot pattern, concentrated in the perinuclear area of myoblasts (Figs. 3A and S6A), and co-localized with thick, well aligned actin stress fibers in both myoblasts (Fig. S6C) and myotubes (Fig. S6I). On the other hand, EGFP-140-kDa palladin appeared as star-like structures (Figs. 3B and S6D). Consistently, myoblasts harboring EGFP-140-kDa palladin (Fig. S6D) exhibited thinner and poorly oriented actin stress fibers, which were also organized in star-like structures (Fig. S6E). The signal of EGFP-200-kDa palladin in myoblasts showed both striated and some star-like structures (Fig. 3C). Interestingly, 140-kDa-palladin-harboring myoblasts had an “unstretched” peripheral plasma membrane (Figs. 3B and S6D), in contrast with that of EGFP-90- and 200-kDa-palladin-expressing myoblasts (Figs. 3A and 3C, respectively). In 4 days-differentiated myotubes, recombinant palladin isoforms also co-localized with both α-actinin (Figs. 3F, 3I and 3L) and actin (Figs. S6I and S6L). However, they showed different myofibril assembly rates. Myotubes expressing EGFP-90-kDa palladin showed the regular expression pattern of α-actinin (Fig. 3E, yellow arrowheads, see inset). However, EGFP-140-kDa palladin failed to form the typical α-actinin sarcomeric pattern (Fig. 3H). Strikingly, the overexpression of EGFP-200-kDa palladin promoted the formation of mature Z-lines even though these myotubes still stayed in early differentiation stage (Fig. 3L, yellow arrows, see inset). Together, the observations suggest palladin isoforms have different roles during myofibril formation.

**Figure 3:**
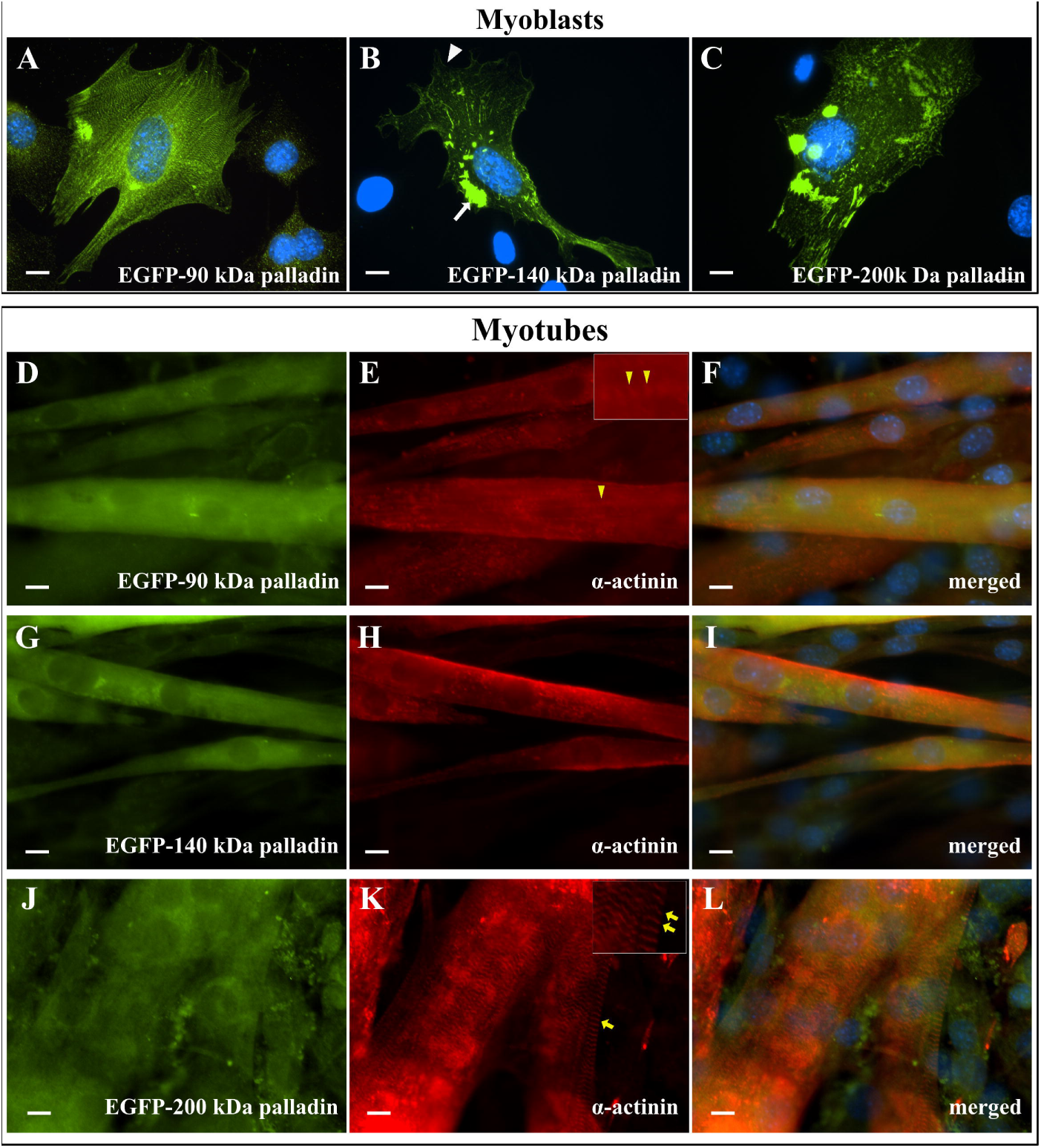
Expression pattern of EGFP-tagged palladin isoforms. Plasmids expressing EGFP-tagged palladin isofroms (green) were used for transfection expresion in C2C12 cells. Myotubes were then immunofluorescence stained with sarcomeric α-actinin (red). Myoblasts expressing (A) EGFP-90-, (B) EGFP-140-, (C) EGFP-200-kDa palladin. (D-F) EGFP-90-kDa palladin expressing myotubes. (G-I) EGFP-140-kDa palladin expressing myotubes. (J-L) EGFP-200-kDa palladin expressing myotubes. Scale bar: 10 μm.

## Discussion

During skeletal muscle differentiation, the cells become elongated and multinucleated, and structures involved in contraction progressively assemble. In premyofibril formation, C2C12 cells reorganize their actin network to become bundles of thin filaments. At the same time, α-actinin becomes more extensively organized into periodic bands along bundles of actin filaments. Subsequently, C2C12 cells rapidly down-regulate the expression of non-muscle myosin and increase the accumulation of adult muscle-specific myosin in the cytoplasm along with other myogenic proteins (Dias et al., 1994; Sabourin and Rudnicki, 2000; Bentzinger et al., 2012). The contractile sarcomeres are organized by the various sarcomeric cytoskeleton proteins, including α-actinin, myomesin, titin, and nebulin. Palladin is known to remodel the actin network and interact with many sarcomeric proteins (Mykkanen et al., 2001; Parast and Otey, 2000; Otey et al., 2005). Thus, it is speculated that palladin is involved in skeletal muscle sarcomere assembly. In this work, the expression pattern of palladin during in vitro myogenesis recapitulated previous reports (Fig. 1A) (Nguyen et al., 2014; Wang and Moser, 2008). The expressions of 90- and 200-kDa palladin were concurrently increased with sarcomeric myosins, suggesting a relationship of palladin with myofibrillogenesis. Our experimental results indicate that palladin is expressed early in myofibrillogenesis. Palladin is co-distributed with major sarcomeric proteins, such as actin, α-actinin, and MyHC, during nascent sarcomere assembly, and then enriched in the I-bands of fully differentiatied myotubes. These findings indicate palladin involvement in the sarcomeric development of cultured myotubes. Present work has led to insights into the interactions and structure of palladin in myofibril assembly.

Sarcomeric α-actinin is an actin-binding protein that cross-links antiparallel actin filaments into mechanically stable structures. α-actinin interacts with palladin and a number of known sarcomeric proteins, such as ALP, ZASP, cypher, myotilin, nebulin, and titin, to maintain ordered myofibril arrays (Pappas et al., 2008; Granzier and Labeit, 2005). It is one of the earliest Z-line markers. Moreover, α-actinin also binds to other membrane-associated proteins, such as vinculin and integrins, to maintain membrane integrity during muscle contraction (Sparrow and Schock, 2009). When differentiation is induced, α-actinin aligns longitudinally across the cytoplasm, cross-links actin filaments, forms a rod-like structure and links all those structures to membrane complexes, giving rise to Z-lines. Palladin is known to co-localize with α-actinin in a variety of cells (Ronty et al., 2004; Beck et al., 2011). In myoblasts, we found that palladin was distributed diffusely with a distinctive beads-on-a-string striated pattern in the cytoplasm or concentrated in focal adhesion sites, co-localized with cytoplasmic actin stress fibers and α-actinin (Fig. 2). In contrast, the typical striated pattern of α-actinin was either not detected or barely detected in the undifferentiated stage. Thus, the appearance of the sarcomeric phenotype of palladin underlines that palladin likely accumulated and became organized into periodic structures independently of α-actinin. Therefore, palladin might be a scaffold for sarcomeric cytoskeletal assembly in skeletal muscle differentiation.

To promote the formation of multinucleated myotubes, differentiating myoblasts activate the expression of the motor protein MyHC. MyHC is one of the main components of the contractile apparatus and acts as a determinant of mature differentiation. MyHC is integrated into the nascent sarcomeres at a late stage of myofibrillogenesis to form the thick filament or A-bands. Our results show that palladin first associates with MyHC in the early myofibril assembly (Fig. S4F), and then presents in the close vicinity of MyHC bands in entirely striated myofibrils (Fig. S4J), when it also co-locolizes with actin and α-actinin of I-bands and Z-lines, respectively. These results indicate that palladin might share some roles with MyHC, which also surrounds the A-bands, in early differentiation. Additionally, our previous work showed that loss of palladin expression in C2C12 nascent myotubes resulted in upregulation of MyHC (Nguyen and Wang, 2015). Thus, it is likely to suggest that MyHC partially compensates for the absence of palladin in this case.

As mentioned earlier, the isoforms of palladin displayed opposite expression patterns during myogenesis - 90- and 200-kDa palladin were increased but 140-kDa palladin was decreased (Fig. 1A). This observation suggests different involvements of the palladin isoforms in myotube maturation. Under differentiation induction, the expression of muscle-specific isoforms is predominantly upregulated and non-muscle-specific isoform expression is concurrently decreased. Therefore, it is reasonable to propose that 90- and 200-kDa palladin, but not 140-kDa palladin, are required for skeletal muscle differentiation. However, isoform-specific palladin antibodies have not yet been commercially developed. Herein, to examine the different roles of palladin isoforms in myogenesis, myoblasts were transfected with EGFP-tagged palladin isoforms and allowed to differentiate into myotubes. The overexpression of 140-kDa palladin in myoblasts bundled the actin network into particular strikingly star-like structures, which is consistent with other reports (Rachlin and Otey, 2006). Since myoblast structure, especially actin cytoskeletons, affects subsequent myotube sarcomere arrangement (Berendse et al., 2003), the expression of α-actinin was examined in palladin-expressing cells. Indeed, 140-kDa-palladin-harboring myotubes were smaller and poorly formed striated α-actinin bands (Fig. 3I). In contrast, 200-kDa palladin overexpression resulted in the building of larger myotubes that are more capable of sarcomere assembly (Fig. 3L). Hence, 200-kDa palladin might facilitate the assembly of sarcomeres. Our observations suggest that palladin isoforms have different roles during myofibril arrangement.

In summary, our results show that palladin displays uneven expression in the C2C12 myoblasts (Fig. 4). Palladin preferentially encircles actin and α-actinin, interconnecting them at the I-band level, forming the scafflods of filaments in sarcomeres. Thus, during sarcomeric assembly, palladin may shortly integrate into the nascent A-bands. On the other hand, palladin associates with I-bands and Z-lines mainly after the formation of mature myofibrils. These findings emphasize the scaffolding role of palladin protein in myofibrillogenesis.

**Figure 4:**
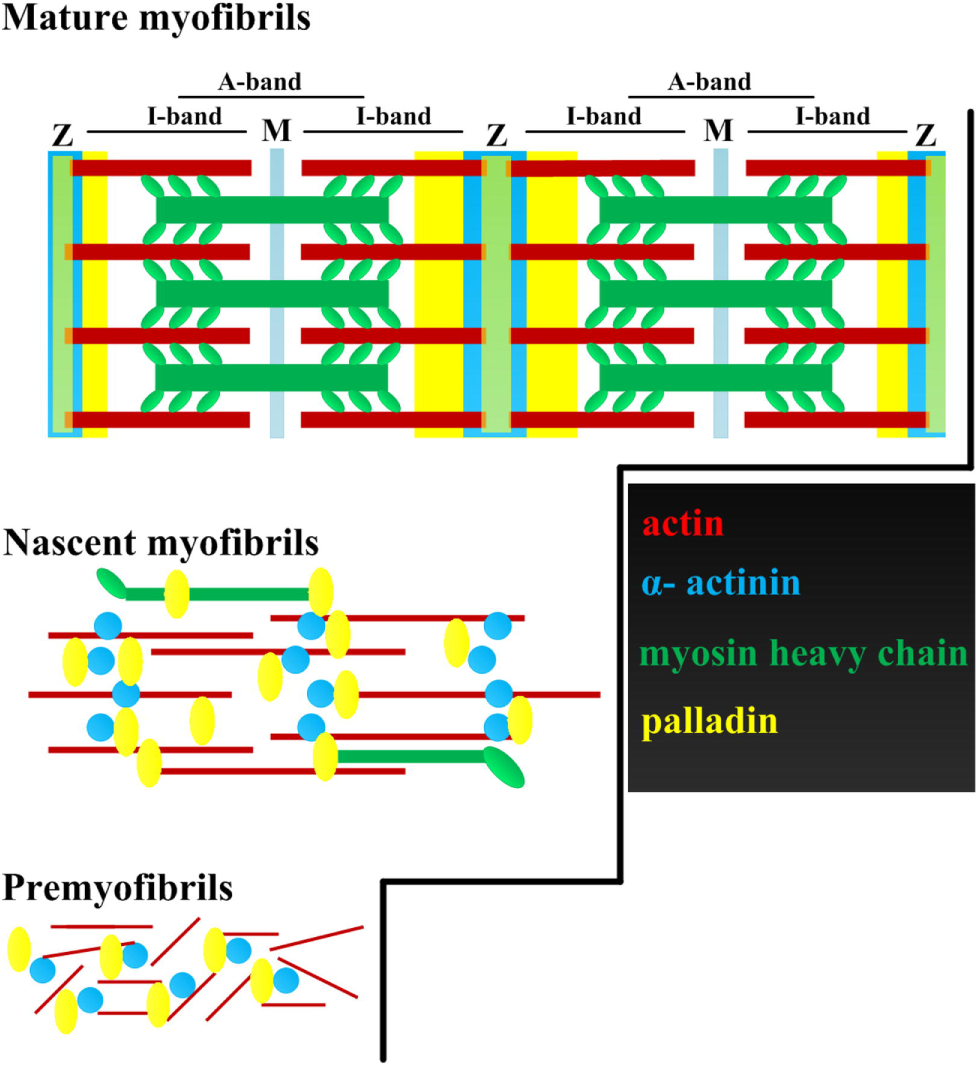
Schematic model outlining the involvement of palladin protein during sarcomeric assembly. Palladin co-localizes with actin and a-actinin during premyofibril. Subsequently, palladin may shortly integrate into the nascent A-bands. However, palladin associates with I-bands and Z-lines mainly during the formation of mature myofibrils.

## Author Contributions

NUNN designed and executed all experiments and data analysis and drafted the manuscript. TYL helped the immunofluorescence preparations. HVW designed the study and reviewed the manuscript. All authors have read and approved the final manuscript.

Supplementary figure legends

Sup Figure 1: Double-immunofluorescence staining of C2C12 myotubes expressing mature myofibrils with sarcomeric α-actinin (red) and myosin light chain (green), imaged by epifluorescence microscopy. Scale bar: 10 μm.

Sup Figure 2: Double-immunofluorescence staining of palladin (green) and actin (red) during myofibrillogenesis, imaged by epifluorescence microscopy. Scale bar: 10 μm.

Sup Figure 3: Double-immunofluorescence staining of palladin (green) and sarcomeric α-actinin (red) during myofibrillogenesis, imaged by epifluorescence microscopy. Scale bar: 10 μm.

Sup Figure 4: Double-immunofluorescence staining of palladin (green) and MyHC (red) during myofibrillogenesis, imaged by epifluorescence microscopy. Scale bar: 10 μm.

Sup Figure 5: Triple-immunofluorescence staining of actin (purple), palladin (yellow), and MyHC (cyan) in adult mouse hind limb skeletal muscle, imaged by epifluorescence microscopy. Scale bar: 5 μm.

Sup Figure 6: Immunofluorescence staining of EGFP-palladin isoforms (green) and actin (red) in overexpressed cells. Scale bar: 10 μm.

